# A marine worm oxygen carrier to improve the quality of preserved tumor samples

**DOI:** 10.64898/2026.11.12.766113

**Authors:** Cottin Maëlle, Bourdages Samuel, Bernèche Alexandre, Ziaee Maryam, Goy Erwan, Péant Benjamin, Kendall-Dupont Jennifer, Cardin Guillaume, Demini Leila, Zal Franck, Christopoulos Apostolos, Gervais Thomas, Yu François, Rodier Francis

## Abstract

Biobanked tissue samples are essential for cancer research, yet pre-analytical ischemia–hypoxia during tissue handling can compromise tissue or derived models’ quality, limiting their value for translational research. Here, we investigated whether M101, an extracellular hemoglobin from *Arenicola marina*, could mitigate indirectly by providing oxygen handling-induced ischemia–hypoxia in cancer tissues and subsequent live 3D models. Using prostate cancer PC3 and ovarian cancer TOV-21G xenografted tumors preserved *ex vivo*, M101 reduced the emergence of hypoxia-associated markers, including the hypoxia-inducible factor 1-alpha, and attenuated metabolic stress. In xenograft-derived micro-dissected 3D tumor models cultured for 8 days, M101 improved tissue morphology, reduced fragmentation, and was associated with higher proliferative activity and lower inflammatory interleukin-6 release. Together, these findings suggest that M101 helps preserve structural and functional features of biobanked tumor models during *ex vivo* handling. Thus, oxygen carriers could represent a bio-compatible strategy to improve the fidelity of biobanked samples and patient-derived tumor models for precision oncology applications.

## 1. Introduction

Biobanks are essential tools in translational oncology, providing preserved tumor tissues but also advanced derived biological material essential to study cancer biology in humans. In particular, biobank-derived models that reflect the disease of origin termed patient “avatars” are essential to explore therapeutic resistance and to advance precision medicine. It is key that both non-living and living biobanks accurately capture molecular and clinical heterogeneity observed across patients, making them central to biomarker discovery and preclinical model development ^1–7^. However, the pre-analytical phase of tissue processing from patients to research laboratories remains a major source of variability. From the time of surgical resection tumor specimens are exposed to warm and cold ischemia, during which blood flow, oxygen and nutrient delivery are abruptly interrupted ^7–10^. This loss of perfusion rapidly induces mitochondrial dysfunction, adenosine triphosphate (ATP) depletion, ionic imbalance and oxidative stress. If oxygen deprivation persists, hypoxia develops and HIF-dependent transcriptional programs are activated. These ischemic and hypoxic conditions can alter metabolism, structure and gene expression, reducing tissue quality, viability and biological representativeness, and ultimately limit the reliability of downstream analyses ^11–19^. Similar concerns apply to patient-derived living models, where suboptimal oxygen conditions during handling and culture may distort functional readouts and drug-response measurements or increase experimental variability, thus reducing analytic power ^20,21^. These challenges, which remain underreported and insufficiently considered, are increasingly important as multi-omics profiling and patient-derived tumor models are used widely in translational research.

In this context, oxygen carriers like the extracellular hemoglobin extracted and purified from the marine annelid *Arenicola marina*, M101, represents an attractive bio-compatible candidate to support oxygen delivery during tissue preservation and *ex vivo* culture. M101 is a giant oligomeric respiratory protein (3600 kDa) with high oxygen-carrying capacity (156 oxygen molecules) and an appropriate oxygen affinity (p50 ≈ 7 mmHg) suited to release oxygen in hypoxic environments ^22–24^. Its stability, supported by an extensive network of disulfide bonds, together with intrinsic cooperativity (n50 ≈ 2.5), a pronounced Bohr effect and an inherent superoxide dismutase–like activity, endows M101 with both oxygen-carrying and antioxidant properties ^25–27^. Formulations based on lugworm extracellular hemoglobins, such as HEMO2life^®^ (*Arenicola marina*) and HEMOXCell^®^ (*Nereis virens*), have been shown to improve tissue oxygenation, reduce ischemia-reperfusion injury and support cell growth in several preclinical and biomanufacturing settings ^26–35^.

Here, we investigated whether M101 could improve the preservation of tumor tissues and living tumor models *ex vivo*. We tested whether supplementing preservation and culture conditions with M101 could mitigate hypoxia and stress-associated alterations while maintaining tissue quality and cellular viability compared with conventional approaches (Fig. 1).

**Fig. 1.**
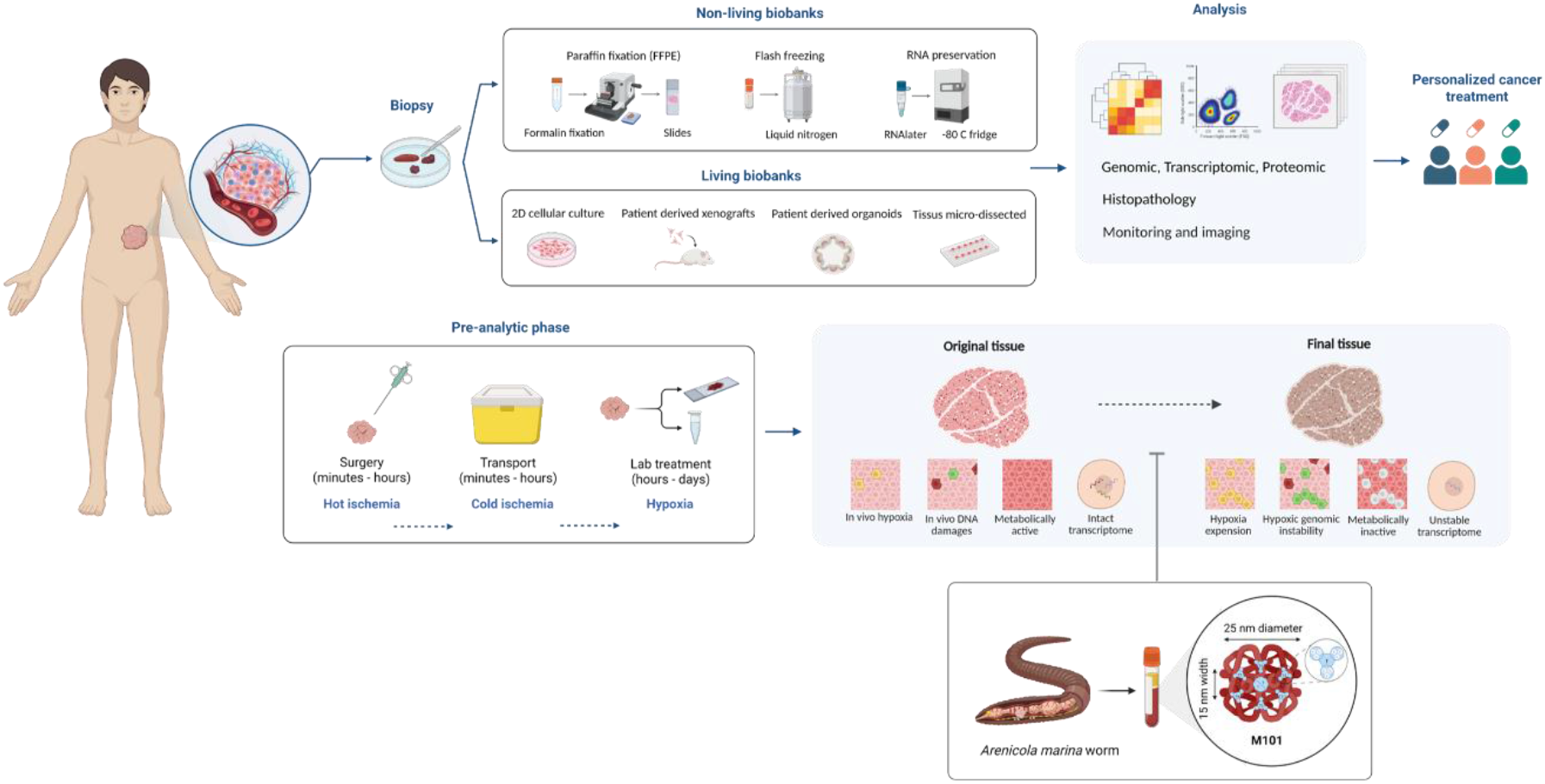
| Overview of pre-analytical ischemia and hypoxia in tumor biobanking and the proposed use of M101. A tumor biopsy is processed into non-living biobanks (formalin-fixed paraffin-embedded tissue, flash-frozen tissue, RNA-preserved samples) or living biobanks (2D cell culture, patient-derived XG, patient-derived organoids, micro-dissected tissue), which support genomic, transcriptomic and proteomic analysis, histopathology, and monitoring and imaging for personalized cancer treatment. The pre-analytical phase comprises surgery (hot ischemia), transport (cold ischemia) and laboratory handling (hypoxia), which can alter the original tissue. The diagram shows M101, the extracellular hemoglobin from Arenicola marina, as a potential additive to be used during the preanalytical phase to counteract these alterations. This figure was created with BioRender.com.

## 2. Results

### 2.1. M101 modulates hypoxia-associated markers in xenograft tissues

To assess the effect of M101 during pre-analytical preservation, we generated prostate cancer PC3 and ovarian cancer TOV-21G xenografts (XG) in NOD-Rag mice. Tumors of approximately 300-400 mm³ were collected and stored on ice in PBS or in PBS supplemented with M101 (1g/L) for 3h30 or overnight (O.N), reproducing a standardized biobanking protocol. Collected tissues were then incubated for 30 min at 37 °C to reveal processes triggered during cold preservation (Fig. 2A,B).

**Fig. 2.**
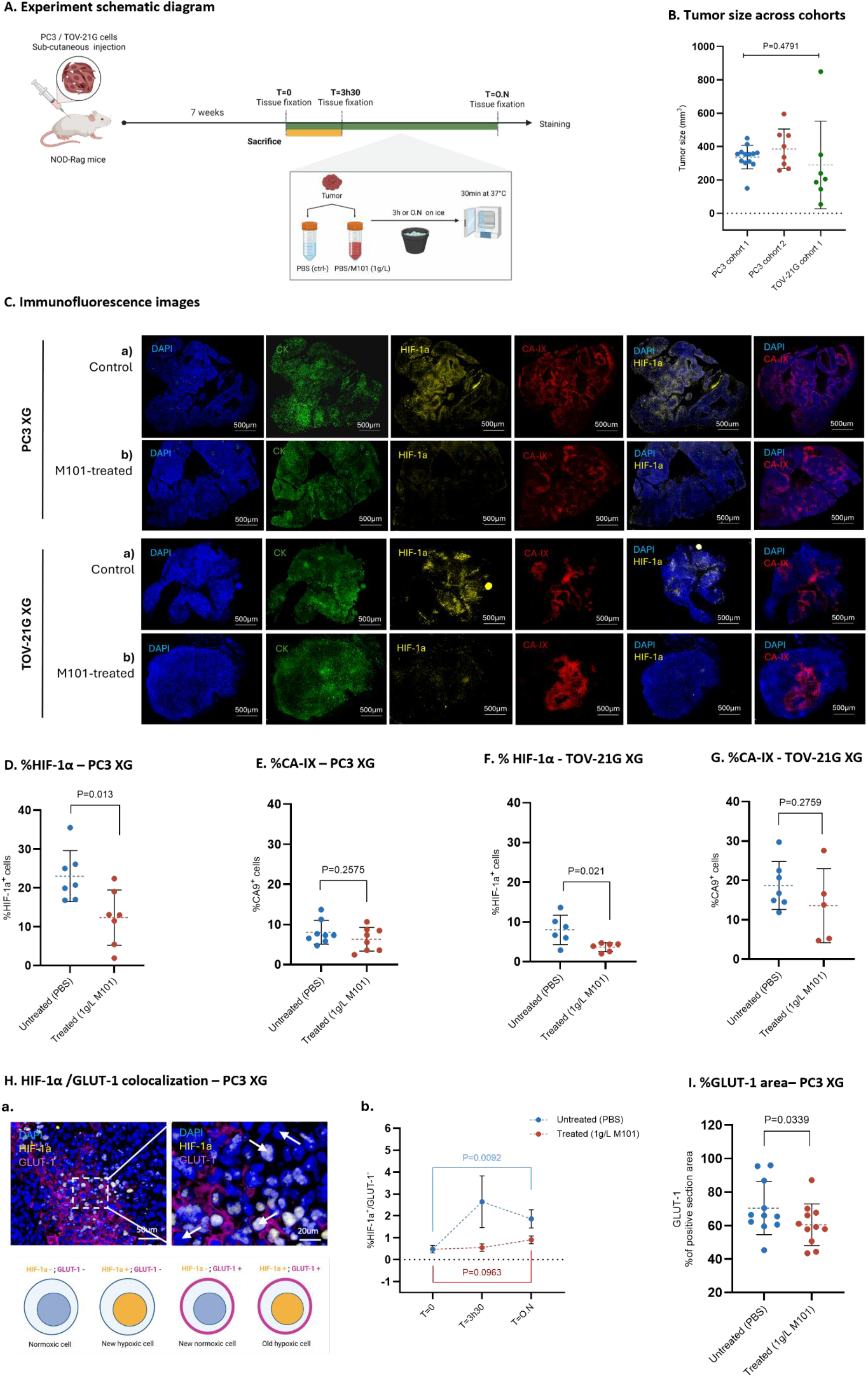
| Effect of oxygen provided by M101 on hypoxia-associated markers in preserved XG tissue. **A.** Experimental workflow showing PC3/TOV-21G XG injection in NOD-Rag mice, tumors recovery and *ex vivo* treatment with PBS or PBS/M101 (1g/L), followed by fixation, RNA collection and staining at the indicated time points. **B.** Tumor size distribution in the two PC3 XG cohorts (n=7; n=13) and TOV-21G XG cohort (n=7). **C.** Representative immunofluorescence images of control (PBS) and M101-treated (1g/L) tissues stained for DAPI, cytokeratin (CK), CA-IX and HIF-1α. Scale bar, 500 µm. **D,E.** Percentage of HIF-1α^+^, and CA-IX^+^ cells in PC3 XG after 3h30 (n=7). **F,G**. Percentage of HIF-1α^+^, and CA-IX^+^ cells in TOV-21G XG after 3h30 (n=7). **H.** (**a)** Representative HIF-1α/GLUT-1 colocalization immunofluorescence and a schematic of the marker combinations in normoxic and hypoxic cells. **(b)** Quantification of HIF-1α^+^/GLUT-1^-^ cells in PC3 XG at T=0, T=3h30 and T=O.N (n=13). **I**. Percentage of GLUT-1^+^ area in PC3 XG after 3h30 (n=13). Data are shown as mean ± SD, with each point representing one tumor. Statistical tests: Unpaired t-test and One-way ANOVA, exact P values are shown in the panels.

Tissues were stained using immunofluorescence for cytokeratin (CK) to reveal epithelial tumor cells, and for hypoxia biomarkers to reveal established hypoxic areas (carbonic anhydrase IX (CA-IX); long half-life) and current hypoxic cells (hypoxia-inducible factor 1-alpha (HIF-1α); short half-life) (Fig. 2C). While all control tumors preserved in PBS-only showed hypoxic areas, as expected for these tumor types ^36^, M101-preserved tumors displayed a significantly reduced proportion of HIF-1α^+^ cells for both PC3 (p=0.013) and TOV-21G (p=0.021) XG (Fig. 2D,F). However, the proportions of CA-IX^+^ cells did not differ between conditions in either model (Fig. 2E,G). This is consistent with the long half-life reported for CA-IX (36–98 h) ^37^, which reflect the pre-existing hypoxic state before tissue collection, while the short half-life HIF-1α probably reflect the actual hypoxic state of cells. To further characterize this dynamic hypoxic state, we examined the ratio of HIF-1α (short half-life) to glucose transporter type 1 ((GLUT-1); (mid half-life)); transcriptional target of HIF-1α that also has a longer half-life ^14,38–42^. In PC3 XG at T = 0, T = 3h30 and T = O.N, the proportions of HIF-1α⁺/GLUT-1⁻ cells (newly hypoxic) increased significantly over time in control tissue (p=0.0092) but not in M101-treated tissue (Fig. 2H). HIF-1α⁺/GLUT-1⁺ cells (hypoxic), HIF-1α⁻/GLUT-1⁺ cells (previously hypoxic) and CA-IX^+^ cells (hypoxic in the last 36 h) did not differ between groups at any time point (Fig. S1 A–C). Given that ischemia–hypoxia alters cellular metabolism during the pre-analytical phase ^41,42^, we next quantified GLUT-1 expression by immunohistochemistry in PC3 tissue. The GLUT-1^+^ area was significantly lower in M101-treated samples (p=0.0339) (Fig. 2I). These metabolic changes were not accompanied by alterations in proliferation (KI67) or cell death (CC3) during the same preservation window (Fig. S1 D-F).

### 2.2. M101 improves the preservation of MDT morphology over time

After assessing non-living biobank fixed tissues, we next examined whether M101 could improve the viability and morphological preservation of 3D living avatars. Fresh PC3-derived XG tissues were mechanically processed to micro-dissected tissues (MDTs), which were then subjected to a 1 h post-sacrifice delay and an additional 30 min handling period before loading into a microfluidic device designed to preserve viability. These specific times were chosen to approximate the average delay between the operating room and the pathology laboratory (1h), and between pathology and the research platform (30min). During the 1h period, tissues were kept dry, as in standard protocols. During the 30min period, tissues were either kept dry (control) or incubated in a solution of HBSS supplemented with M101 (1g/L). After punching and loading, MDTs were cultured for 8 days with or without M101 (1g/L). (Fig. 3A,B).

**Fig. 3.**
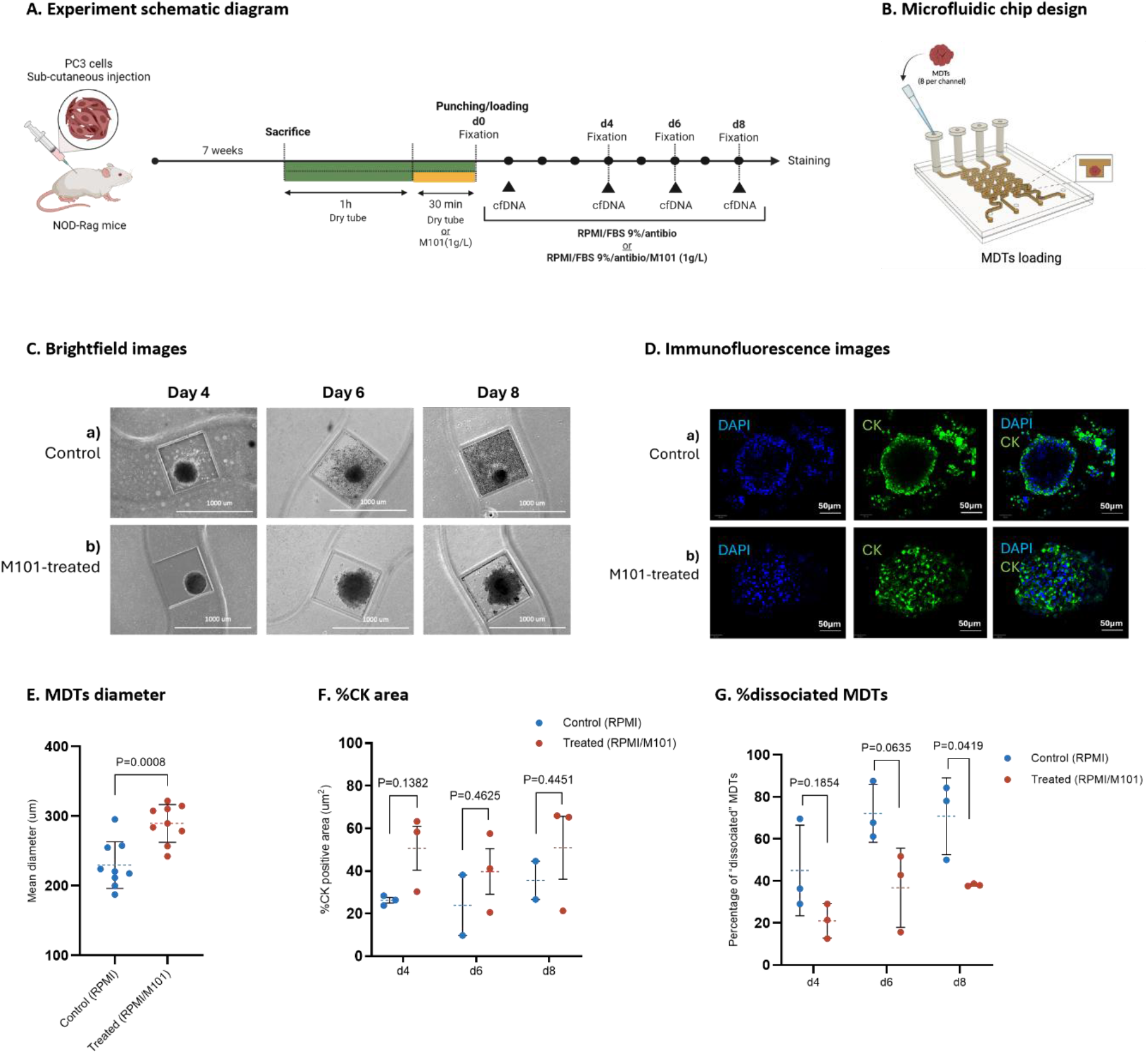
| Effect of M101 on the morphology of MDTs over time. **A.** Experimental timeline from PC3 XG generation to MDT production, loading and culture under control (RPMI) conditions or RPMI supplemented with M101 (1g/L), followed by samples collection and staining at the indicated time points. **B.** Schematic of the microfluidic chip used for MDT loading and culture. **C.** Representative brightfield images of control and treated MDTs at d4, d6 and d8. Scale bar, 1000 µm. **D.** Representative immunofluorescence images of control and M101-treated (1g/L) MDTs stained for CK. **E.** Mean diameter of control and M101-treated (1g/L) MDTs. **F.** CK^+^ area over time. **G.** Percentage of dissociated control and M101-treated (1g/L) MDTs over time. Data are shown as mean ± SD, with each point representing one independent experiment. Statistical tests: Unpaired t-test, exact P values are shown in the panels.

Overall, M101-treated MDTs exhibited less pronounced morphological alterations and retained larger tissue structure at day 8. Indeed, brightfield and CK imaging showed progressive fragmentation of control MDTs over time (Fig. 3C,D), consistent with the observed decrease in mean diameter and the rise in the proportion of dissociated MDTs (Fig.3E-G).

### 2.3. M101 enhances MDTs proliferative capacity over time

We next examined whether the improved structural preservation of M101-treated MDTs was associated with changes in hypoxia, stress and viability during culture, despite the small size of these models, designed to limit the formation of a hypoxic core (Fig. 4A,B) ^43^. Indeed, both HIF-1α and CA-IX signal remained low in both conditions throughout the 8-day culture period (Fig. 4C,D). To determine whether M101 affected cell dynamics during culture, we evaluated proliferation and cell death markers. The proportion of KI67^+^ cells differed between conditions: M101-treated MDTs showed higher KI67 positivity that remained closer to the baseline (p=ns) than control MDTs (p=0.0030) (Fig. 4E). Regarding apoptosis rate, the proportion of CC3^+^ cells showed an overall increase at day 4, followed by a slight decrease at days 6 and 8, without significant differences between conditions (Fig. 4F). Quantification of cfDNA concentration in the culture supernatant, as a complementary readout of cell death ^44^, showed a similar temporal pattern, with higher concentrations at earlier time points and lower levels by day 8 (Fig. 4G). We also assessed whether M101 influenced cellular stress by quantifying interleukin-6 (IL-6) concentration in the cultured medium. IL-6 levels were elevated during the first 4 days of culture in control MDTs, while M101-treated MDTs released lower amounts from day 1 onwards and maintained relatively stable levels over the culture period (Fig. 4H).

**Fig. 4.**
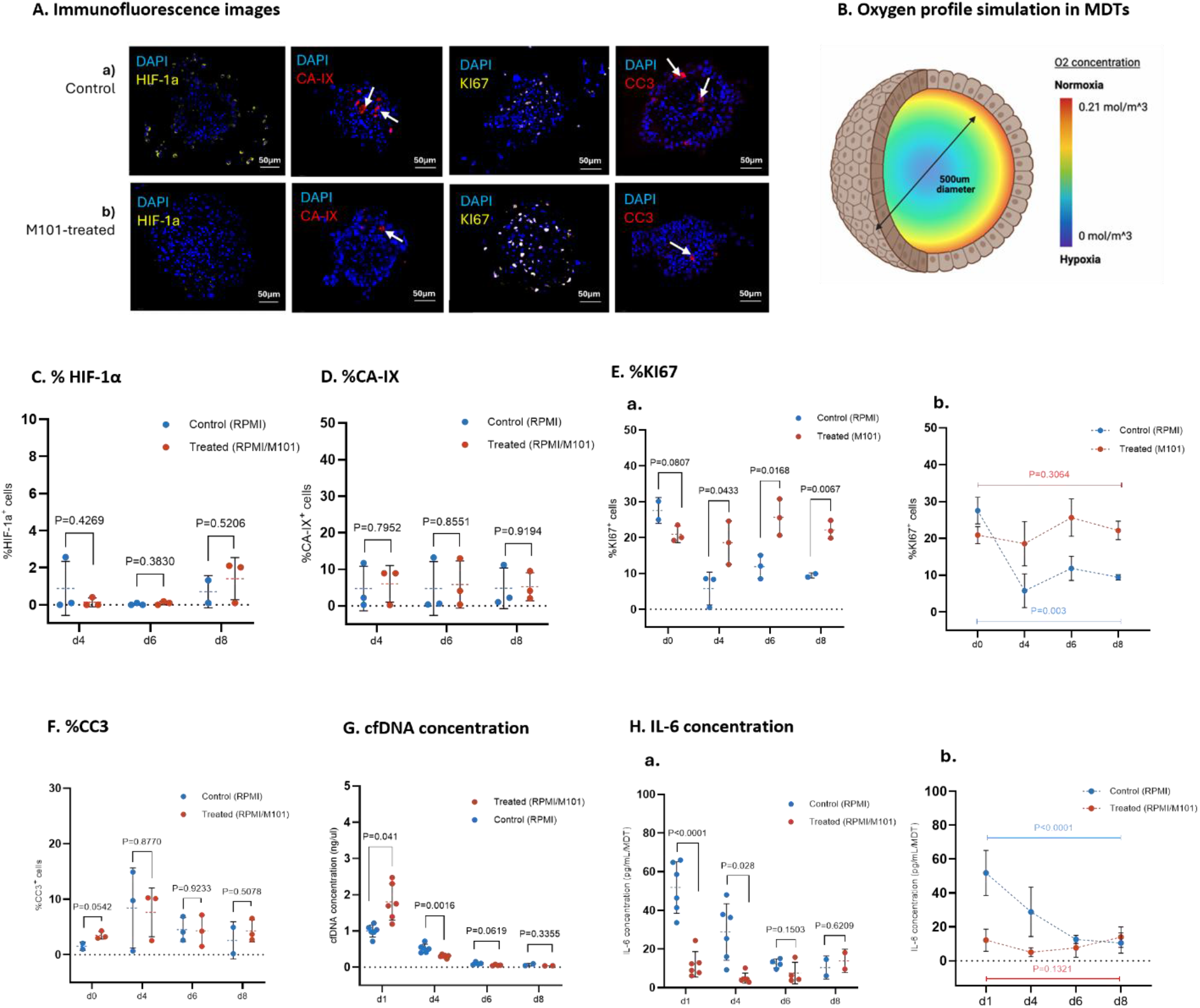
| Effect of M101 on proliferation, cell death and IL-6 release in cultured MDTs. **A.** Representative immunofluorescence of control (RPMI) and M101-treated (1g/L) MDTs stained for DAPI, CA-IX, HIF-1α, KI67 and CC3. Scale bar, 50 µm. **B.** Schematic and simulated radial O2 profile for 500 µm diameter MDTs. **C,D.** Percentage of HIF-1α⁺ and CA-IX⁺ cells over time. **E.** Percentage of KI67⁺ cells over time. **F.** Percentage of CC3⁺ cells over time. **G.** Quantification of the total cfDNA concentration in supernatant over time (ng/µl). **H.** Quantification of IL-6 concentration in culture supernatant over time (pg/mL). Data are shown as mean ± SD, with each point representing one independent experiment, or mean value for combined MDTs for each condition. Statistical tests: Unpaired t-test and One-way ANOVA, exact P values are shown in the panels.

## 3. Materials and methods

### 3.3. Cell lines

Human prostate cancer cell line PC3 (ATCC) and human ovarian cancer cell line TOV-21G (provided by Dr. Anne-Marie Mes-Masson) were respectively maintained as monolayers in RPMI (350-000-CL,Wisent,Saint-Jean-Baptiste, QC,Canada) and OSE (316-030 CL Wisent,Saint-Jean-Baptiste, QC,Canada) medium, both supplemented with 9% fetal bovine serum (FBS; Gibco, Thermo Fisher, Waltham, MA, USA), 100 IU/mL penicillin and 100 µg/mL streptomycin (450-201-EL, Wisent, Saint-Jean-Baptiste, QC, Canada), and maintained at 37 °C in 20% O_2_ and 5% CO_2_ conditions. Medium was replaced every 2–3 days, and cells were passaged at approximately 80% confluence. Cultures were routinely tested for mycoplasma contamination.

### 3.4. Mouse models

Male immunodeficient NOD-Rag mice (NOD.Cg-*Rag1^tm1Mom^ Il2rg^tm1Wjl^*/SzJ, The Jackson Laboratory, Canada, stock #007799; RRID:IMSR_JAX:007799) ^45^ were used to generate PC3 XG. Female immunodeficient NOD-Rag mice (NOD.Cg-*Rag1^tm1Mom^ Il2rg^tm1Wjl^*/SzJ, The Jackson Laboratory, Canada, stock #007799; RRID:IMSR_JAX:007799) were used to generate TOV-21G XG. All animal procedures were performed in accordance with institutional and national guidelines for the care and use of laboratory animals (Ethic committee approval number: I23040FYs (PC3); C23030FRs (TOV-21G)).

### 3.5. M101

The extracellular hemoglobin M101 (HEMARINA, Morlaix, France), derived from *Arenicola marina*, was provided as a 20 mL vial (50.5 g/L) and was aliquoted into 500 µL fractions (1g/L working concentration) or 300 µL fractions (50.5 g/L stock) and stored at −80 °C. 1g/L concentrations were used for biobank sample preservation and *ex vivo* micro-dissected tissue (MDT) culture ^46^.

### 3.6. XG generation

PC3 and TOV-21G cells were harvested by trypsinization (0.025% trypsin/0.221 mM EDTA (325-542-EL, Wisent, Saint-Jean-Baptiste, QC,Canada), 5 min at 37 °C), neutralized, counted using a TC20 Automated Cell Counter (Bio-Rad), and centrifuged at 600×g for 5 min. Cell pellets were resuspended at 2×10^7^ cells/mL in the appropriate culture medium. For biobank XG, 50 µL of cell suspension was mixed 1:1 with Matrigel (Corning, USA) and injected bilaterally in the subcutaneous flank. Tumors were harvested when they reached approximately 350 mm^3^. For MDTs production, 50 µL of the same suspension was injected unilaterally, and tumors were excised at a mean volume of approximately 1000 mm^3^. The two PC3 XG cohorts had respectively n=7 n=13 tumors. The TOV-21G XG cohort had n=7 tumors.

### 3.7. Tissue collection and preservation

Excised XG tumors were aseptically sectioned into equal parts corresponding to the different fixation time points. To mimic pre-analytical delay, samples were stored on ice (4°C) in Dulbecco’s Phosphate-Buffered Saline (PBS (311-425 LL, Wisent, Saint-Jean-Baptiste, QC,Canada)) or in PBS supplemented with M101 (1g/L) for 3h or overnight (O.N). Samples were then incubated at 37 °C (in an incubator) for 30 min to reveal cellular or molecular phenotypes induced during cold preservation. Tissues were directly fixed in 10% formalin (#F-150, ChapTec, Canada) for 24h, or stored in RNAlater (Invitrogen, Thermo Fisher Scientific, AM7021) for downstream analysis.

### 3.8. Microfluidic devices pre-treatment

Microfluidic chips (MISOchip®) consisted of two polydimethylsiloxane (PDMS, Sylgard® 184 silicone elastomer kit, Dow Corning, Midland, USA) layers molded from polymethyl methacrylate (PMMA) or 3D-printed resin masters. The lower layer contained four rectangular fluidic channels (0.9 × 1.1 mm cross-section) with individual inlets (3.2 mm of diameter) and outlets (1.5 mm of diameter). Each channel overlaid a grid of 4 × 8 square wells (0.7 mm) used to trap MDTs. Device surfaces were treated O.N with Pluronic® F-108 (Sigma-Aldrich, St. Louis, USA) to prevent cell adhesion to PDMS. Air bubbles were removed using 100% ethanol, then devices were sterilized with 70% ethanol for 15 minutes. The channels were rinsed with sterile Hanks’ Balanced Salt Solution (HBSS, #311-516-CL, Wisent Inc., Saint-Bruno-de-Montarville, Canada) and placed in a humidified cell incubator (37 °C, 5% CO2) ^47^.

### 3.9. MDT generation and culture

After tumor excision, samples were divided into two equal pieces. One fragment was left dry for 1 h to simulate surgical delay and then incubated for 30 min in a solution of HBSS/M101 (1g/L), whereas the matched fragment was kept dry as a control. Tumors were sliced into 350 µm sections using a tissue chopper (McIlwain, Ted Pella, CA, USA) and punched into 500 µm MDTs preserved in HBSS supplemented with 10% FBS, gentamicin (55 mg/L) (Wisent Inc.) and amphotericin B (0.6 mg/L) (Wisent Inc.). Eight MDTs were loaded manually into each channel (32 MDTs per device). HBSS was then replaced with culture medium (RPMI/FBS/gentamicin/amphotericin B). Medium was exchanged 24h after loading and every 48h thereafter. Control chips received standard medium, whereas treated chips received medium supplemented with M101 (1g/L). MDTs were cultured for 8 days.

### 3.10. Cell-free DNA (cfDNA) quantification

Culture media from MDTs were collected on days 1, 4, 6 and 8. Samples (approximately 60 µL per channel) were centrifuged at 1,500 rpm for 5 min at 4°C, and 10 µL of supernatant was frozen at −80 °C. cfDNA was quantified by quantitative PCR (qPCR) using 10 ul mix of PowerTrack SYBR Green Master Mix (Invitrogen, #11762-500), 50 µM forward and reverse primers targeting human long interspersed nuclear element (LINE) sequences (Fwd: 5’-TCACTCAAAGCCGCTCAACTAC-3’; Rev: 5’-TCTGCCTTCATTTCGTTATGTACC-3’) (generated by Invitrogen, ThermoFisher Scientific, Ontario, Canada), nuclease free water, and media samples in 10:1:8 (v/v ratio). Samples were prepared into MicroAmp Fast Optical 96-Well Reaction Plate (ThermoFisher Scientific) and run in Bio-Rad CFX96 Real-Time System. Standard curves were generated using serial dilutions of mouse genomic DNA (0.0001–100 ng/µL) (69239, MilliporeSigma), and cfDNA concentrations were calculated accordingly.

### 3.11. IL-6 quantification

IL-6 levels were measured in MDTs culture supernatants collected on days 1, 4, 6 and 8 using 96-well ELISA plates (R&D Systems, DY990) according to the manufacturer’s instructions. Optical density was measured at 450 nm with 570 nm background correction.

### 3.12. Fixation and paraffin embedding

#### a. Biobank tissues

Tissue fragments were fixed in 10% formalin for 24 h, transferred to 70% ethanol and processed for paraffin embedding at the CHUM platform. Blocks were sectioned at 4 µm, mounted on TOMO hydrophilic adhesion slides (Matsunami) and stained with hematoxylin and eosin (H&E).

#### b. MDTs

MDTs were fixed in formalin with two 5-min rinses followed by a 30-min incubation. Samples were stained with phloxine B for 20 min, dehydrated through graded ethanol (50–100%), cleared with xylene substitute and embedded in Paraplast X-Tra (Sigma-Aldrich). Sections (4 µm) were mounted on TOMO hydrophilic adhesion slides and stained with H&E.

### 3.13. Immunofluorescence and immunohistochemistry

#### a. Antibodies

The following markers were analyzed: DAPI for nuclei; cytokeratin (CK) 7/18/19 for epithelial tumor cells; HIF-1α and carbonic anhydrase IX (CA-IX) for hypoxia; glucose transporter 1 (GLUT-1) for metabolism; KI67 for proliferation; cleaved caspase-3 (CC3) for cell death; 4-hydroxynonenal (4-HNE) for oxidative stress; phosphorylated histone H2AX (γ-H2AX) for DNA damage.

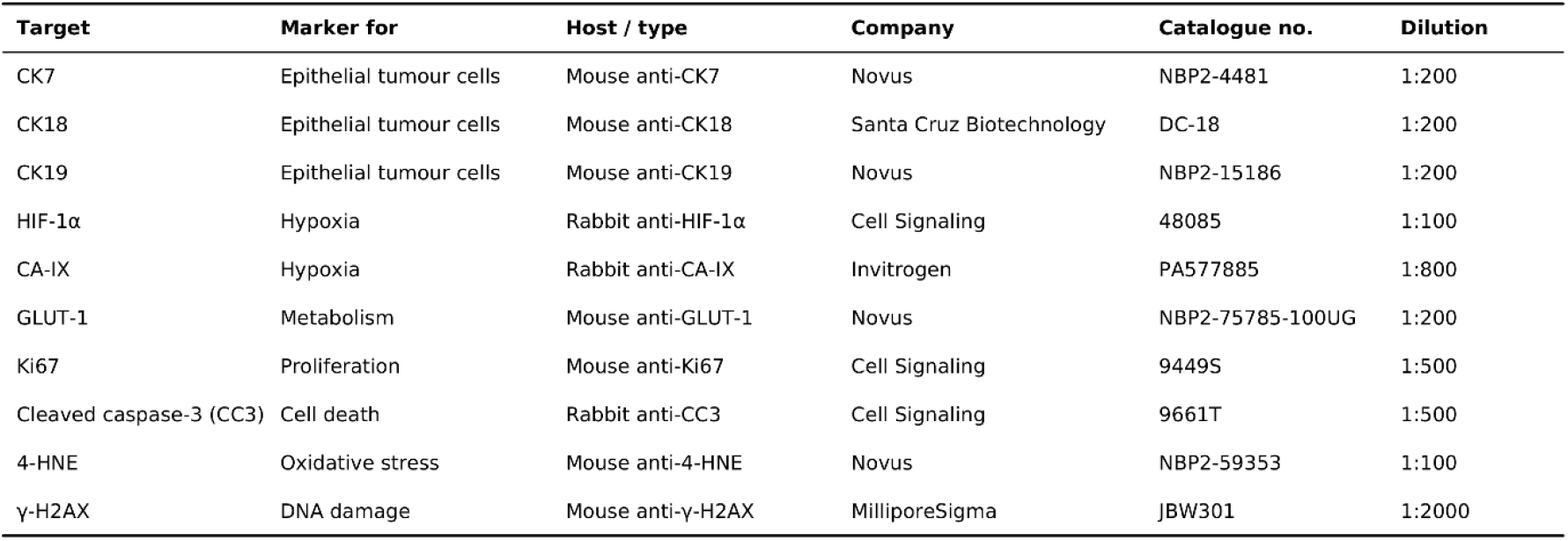

| Target | Marker for | Host / type | Company | Catalogue no. | Dilution |
|---|---|---|---|---|---|
| CK7 | Epithelial tumour cells | Mouse anti-CK7 | Novus | NBP2-4481 | 1:200 |
| CK18 | Epithelial tumour cells | Mouse anti-CK18 | Santa Cruz Biotechnology | DC-18 | 1:200 |
| CK19 | Epithelial tumour cells | Mouse anti-CK19 | Novus | NBP2-15186 | 1:200 |
| HIF-1α | Hypoxia | Rabbit anti-HIF-1α | Cell Signaling | 48085 | 1:100 |
| CA-IX | Hypoxia | Rabbit anti-CA-IX | Invitrogen | PA577885 | 1:800 |
| GLUT-1 | Metabolism | Mouse anti-GLUT-1 | Novus | NBP2-75785-100UG | 1:200 |
| Ki67 | Proliferation | Mouse anti-Ki67 | Cell Signaling | 9449S | 1:500 |
| Cleaved caspase-3 (CC3) | Cell death | Rabbit anti-CC3 | Cell Signaling | 9661T | 1:500 |
| 4-HNE | Oxidative stress | Mouse anti-4-HNE | Novus | NBP2-59353 | 1:100 |
| γ-H2AX | DNA damage | Mouse anti-γ-H2AX | MilliporeSigma | JBW301 | 1:2000 |

#### b. Staining

Formalin-fixed paraffin-embedded (FFPE) tissue and MDT sections were processed on a Discovery Ultra Ventana platform (Roche) using Tris-EDTA antigen retrieval (pH 7.8). Slides were incubated with primary antibodies for 60 min at 37 °C and detected using fluorescent or HRP-based systems. Slides were scanned on an Aperio Versa 200 scanner (Leica Biosystems) and viewed using Aperio ImageScope^®^.

#### c. Image quantification

Protein expression by immunofluorescence (IF) or immunohistochemistry (IHC) was quantified using the open-source QuPath^®^ software (v.0.6.0). Necrotic areas and artifacts were excluded. DAPI^+^ nuclei defined total cell counts, and CK staining delineated epithelial tumor cells. HIF-1α, CA-IX and KI67 were quantified as the percentage of positive tumor cells. NF-κB, CC3 and 4-HNE were measured as mean fluorescence intensity (MFI), and γ-H2AX foci were counted per nucleus. In IHC analysis, GLUT-1 was quantified as the ratio of positive pixel area to total tumor area.

### 3.14. RNA extraction and analysis

Total RNA was extracted from frozen tumor samples in RNAlater, homogenized in Trizol. After chloroform phase separation, RNA was precipitated with isopropanol, washed with 75% ethanol and resuspended in RNase-free water. RNA quantity and purity were measured using a NanoDrop spectrophotometer, and RNA integrity was assessed on an Agilent Bioanalyzer 2100 using an RNA 6000 kit. RNA integrity number equivalent (RINe) value >6 was considered as high quality.

### 3.15. Statistical analysis

Statistical analyses were performed using GraphPad Prism^®^ v9.5.0. At least 5 tissue sections per time point and condition were analyzed. For *in vivo* studies, n refers to mice number. For *ex vivo* studies, n refers to independent experiments. Data are presented as mean ± SD. Statistical significance was set at p < 0.05 using Student’s t-test, One-way or two-way ANOVA, as appropriate and specified.

## 4. Discussion

Optimizing tumor tissue preservation remains a central challenge in translational oncology, where biopsy and resection specimens underlie individualized therapeutic modeling and the development of personalized treatment strategies. Numerous studies have shown that ischemic–hypoxic stress arising during tissue collection, transport and storage can rapidly induce major cellular and molecular alterations. Historically, efforts to mitigate this stress have focused on improving rapid-cooling protocols and shortening the pre-analytical interval ^7,48,49^. More recently, oxygenated microfluidic devices have enabled better control of tissue oxygenation during *ex vivo* culture ^43^. However, no innovative oxygen-delivering agent has yet been tested to continuously restore and maintain adequate oxygen levels during *ex vivo* latency phases or to compensate for uncontrolled spatial and temporal variations in oxygen tension within the tissue.

Our study addresses this issue by evaluating whether M101, a marine hemoglobin with oxygen-carrying capacity, could limit these changes and be used as a simple additive for *ex vivo* preservation of tumor tissues and living micro-dissected tumor models. Across these complementary models, oxygen provided by M101, when used during the warming of tissue from 4°C to 37°C, was associated with reduced early hypoxia signalling and altered expression of hypoxia-associated metabolic markers, improved MDTs structure and lower early IL-6 release. Together, these findings support a model in which M101 helps buffer early handling-related stress rather than reversing an established hypoxic state.

In PC3 and TOV-21G XG preserved under standardized biobanking conditions, M101 reduced the emergence of HIF-1α^+^ cells during the thermal transition from ice bath to the incubator. This effect was accompanied by a reduction in GLUT-1^+^ area in PC3 tissues, consistent with a lower requirement for hypoxia-associated glucose uptake. CA-IX was unchanged, consistent with its long half-life, which suggests that M101 acts on the early, dynamic phase of the hypoxic response rather than on established hypoxic programs. Longer time-course experiments (over several days) will be needed to determine whether M101 can also reduce CA-IX levels. The concordant changes in HIF-1α and GLUT-1 have a plausible mechanistic basis. HIF-1α is rapidly stabilized when oxygen is low and drives transcription of glycolytic genes, including *SLC2A1* (GLUT-1). Limiting the emergence of HIF-1α⁺ cells would therefore be expected to dampen downstream GLUT-1 induction. When cells go from 4°C to 37°C, the solubility of O_2_ in the tissue decreases by a factor of 2. Simultaneously, O_2_ consumption via the accelerated metabolism increases ∼40-fold ^50^. This double effect suppresses hypoxia entirely in cold cells. At 1g/L, the presence of M101 increases the total amount of O_2_ concentration in the tissue by 21% (0.24 mM) with respect to the total dissolved O_2_ in the medium (0.2 mM) at 37°C. As cells warm up, it is hypothesized that the O_2_ is released from the macromolecules as the cellular consumption increases, effectively postponing the onset of hypoxia over the 30 min of the temperature transient. Once the temperature has stabilized, M101 reaches equilibrium within the surrounding hypoxic regions. However, as we did not measure oxygen tension directly, this interpretation is inferential. These hypoxic and metabolic changes were not accompanied by detectable differences in proliferation or apoptosis in this model as both groups showed expected gradual decline in proliferative activity during cold preservation. This dissociation between metabolic marker expression and cell-fate readouts is not surprising as proliferation and apoptosis are comparatively slow, energy-intensive processes, whereas the transcriptional and post-translational changes captured by HIF-1α and GLUT-1 can occur within minutes to hours. Longer delays or more sensitive assays may be needed to test those cell fates.

A key concern for translational use is whether M101 introduces unintended molecular damage, for example through excessive oxygen delivery. We found no increase in 4-HNE, a marker of lipid peroxidation and oxidative stress, nor in γ-H2AX, which correspond to the phosphorylated form of histone H2AX that forms nuclear foci at sites of DNA double-strand breaks ^51^. Neither marker increased in any of the metrics examined (Fig. S2A-D). These findings show that, under these conditions, M101 does not induce detectable oxidative or genotoxic stress, supporting its compatibility with standardized biobanking workflows.

Because ischemia–hypoxia can affect RNA quality from approximately 4h of preservation onwards ^52^, we assessed RNA integrity at T = 0, T = 3h30 and T = O.N RINe values were relatively high (>6) in both groups, and no difference was detected between control and M101-treated tissue at any time point (Fig. S1G,). This result indicates that our preservation window was insufficient to produce a measurable global loss of RNA quality in either condition, or that M101 did not adversely affect the transcriptome over this interval. A more sensitive assessment of transcriptomic changes, including bulk RNA sequencing, could clarify whether M101 influences specific gene expression programs beyond what is captured by RINe alone.

In living MDTs, M101 was associated with better conservation of morphology, reduced fragmentation and larger retained tissue structures over 8 days of culture. Cytokeratin staining further indicated more preserved epithelial organization in treated samples. These data are consistent with a model in which reduced early stress signalling helps maintain tissue cohesion, rather than simply preventing overt cell death. This interpretation is supported by the persistence of proliferative activity in treated MDTs, which remained closer to baseline than in controls, and by the lower IL-6 release observed during the first days of culture. Because the first 24–48 hours after tissue processing are typically the most stressful, this effect may be especially relevant for preserving samples intended for functional assays or drug testing. Overall, the observed effects plausibly reflects buffering of the ischemic and handling stress at the transition from *in vivo* to *ex vivo* conditions. In this sense, M101 could complement, rather than replace, rapid cooling and microfluidic oxygenation.

Several limitations should be considered. First, most experiments were mainly performed in specific tumor model, which restricts generalization to other tumor types with different vascular, metabolic or stromal features, so validation in additional models and patient-derived samples would be needed. Furthermore, because M101 is poorly immunogenic and no specific antibody could be developed against it, its distribution within tissues has not been measured directly, leaving its penetration and spatial exposure uncertain. The link between M101 and the reduced hypoxia-associated signalling (HIF-1α, GLUT-1) is therefore inferred from indirect markers. Functional metabolic assays, such as glucose uptake and oxygen consumption, are also needed to confirm the GLUT-1 findings.

MDTs models also have their specific limitations. The model is derived from XG tissue rather than patient samples, and microdissection, slicing, punching and manual loading may introduce variation in fragment size, cellular composition and tissue integrity. Regarding the morphological findings, larger fragments could also be affected by altered compaction, fixation or imaging properties. Finally, the translational relevance of M101 remains to be established. The current study evaluated a single concentration and did not define a dose–response relationship, the optimal exposure duration or the persistence of M101 during storage and culture. Potential effects on downstream applications—including immunostaining, RNA sequencing, genomic analyses and drug-response assays—also require systematic evaluation. Bulk RNA sequencing may identify transcriptomic changes that are not reflected by RINe, but such changes will need to be interpreted alongside functional data to distinguish beneficial preservation from treatment-induced biological alteration.

Overall, our findings identify M101 as a promising additive for *ex vivo* tumor preservation, and the present study should be viewed as a proof of concept. M101 attenuated early hypoxia-associated responses during thermal transitions and was compatible with the short-term workflows tested. In living MDTs, it helped maintain tissue architecture and proliferative competence. Overall, it also did not affect the tissue survival or displayed any toxicity. These properties could be useful for biobanking, tumor modeling and preclinical drug testing, where sample integrity is essential for reliable interpretation. However, our data do not yet establish direct tissue oxygenation, universal applicability or improved performance of clinical models. Addressing these questions in patient-derived material, with direct oxygen and functional measurements, should guide the next phase of validation. More broadly, this work supports the idea that controlled oxygen delivery may improve the quality and functional relevance of tumor specimens handled outside the body.

## Acknowledgements

We thank Audrey Glory, Kim Leclerc Desaulniers and the staff of the animal facility of the Centre de recherche du Centre hospitalier de l’Université de Montréal (CRCHUM) for support with mouse care, tumor injection and measurements. We thank Véronique Barres and Liliane Meunier of the Molecular Pathology core facility of the CRCHUM for their advice and for performing paraffin sectioning, automated immunofluorescence staining on the Ventana platform and slide scanning. We thank Amélie St-Georges-Robillard of the Microfluidic core facility of the CRCHUM for their advice and assistance with MDT production and analysis. This work was funded by the Québec Consortium for Drug Discovery (CQDM) and the Dr. Azar-Angelil Research Chair in ENT Oncology at the University of Montreal (UdeM). This research was also supported by the Institut du cancer de Montréal (A.C, T.G, F.Y and F.R), the Canadian Institute for Health Research (CIHR MOP114962 to F.R.), the Cancer Research Society (CRS) in partnership with Ovarian Cancer Canada (22713 to F.R.), the Natural Sciences and Engineering Research Council of Canada (NSERC) (F.R.), the Terry Fox Research Institute (TFRI) (T.G., F.Y. and F.R.) and by the Fonds de recherche du Québec – Santé (FRQS) réseau cancer (RRCancer). A.C, T.G., F.Y. and F.R. are researchers of CRCHUM/ICM, which receive support from the FRQS. M.C. and S.B. were supported by MITACS scholarships and by Canderel studentship from the Institut du cancer de Montréal.

