## Supplemental figures for "A marine worm oxygen carrier to improve the quality of preserved tumor samples"

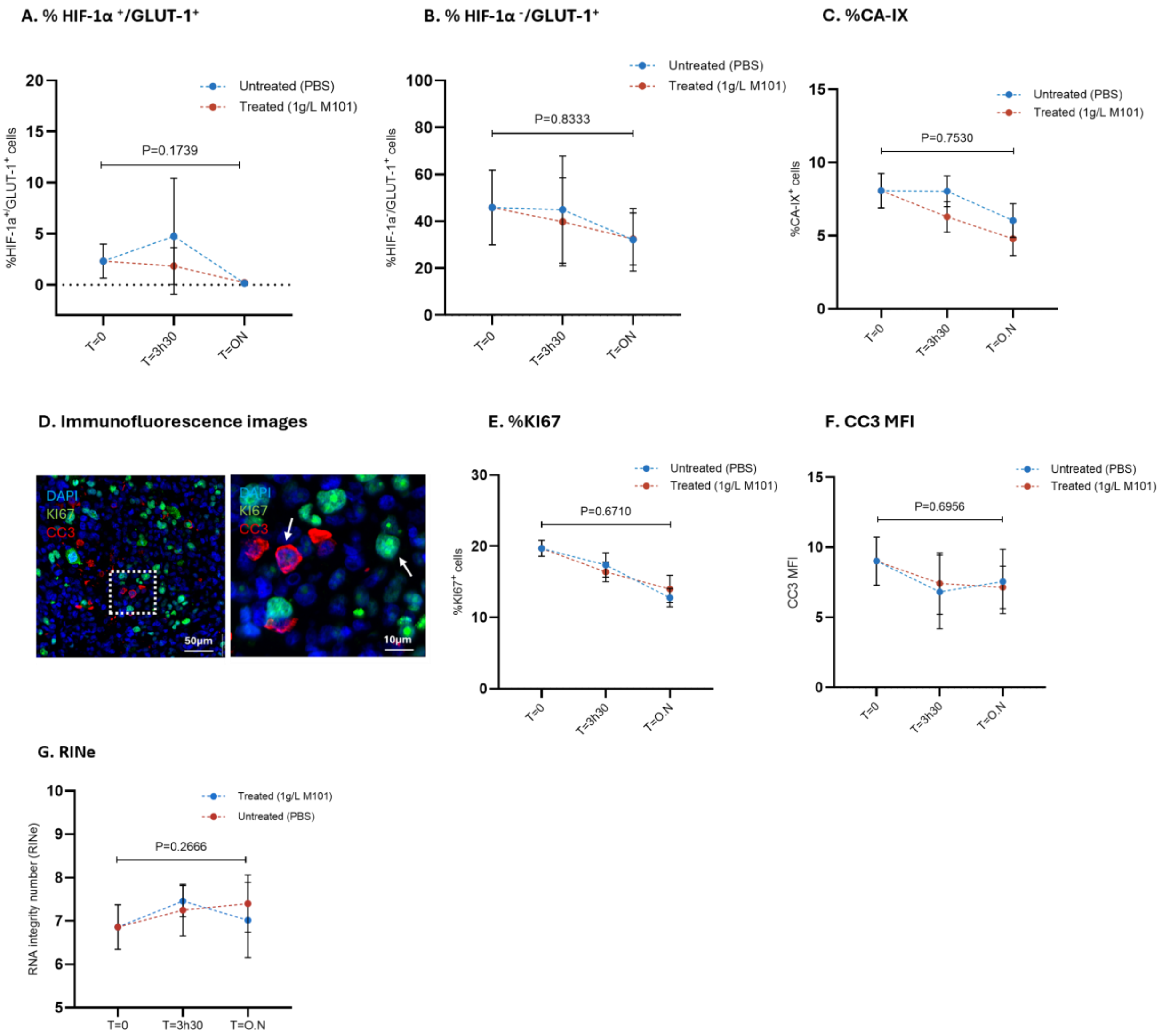


**Fig. S1 | Additional marker analysis in preserved PC3 XG tissue.**

**A,B**. Percentage of $HIF\text{-}1\alpha^+/GLUT\text{-}1^+$ cells and $HIF\text{-}1\alpha^-/GLUT\text{-}1^+$ cells, at T = 0, T = 3h30 and T = O.N in control (PBS) and M101-treated (1g/L) tissue. **C.** Percentage of $CA\text{-}IX^+$ cells over time. **D.** Representative immunofluorescence images stained for DAPI, CK, KI67 and CC3. Scale bar, 50 µm and 10 µm **E.** Percentage of $KI67^+$ cells over time. **F.** CC3 MFI over time. **G.** RNA integrity number equivalent (RINe) over time. Data are shown as mean ± SD, with each point representing one tumor, or the mean value for combined tumors. Statistical tests: Two-way ANOVA, exact P values are shown in the panels.

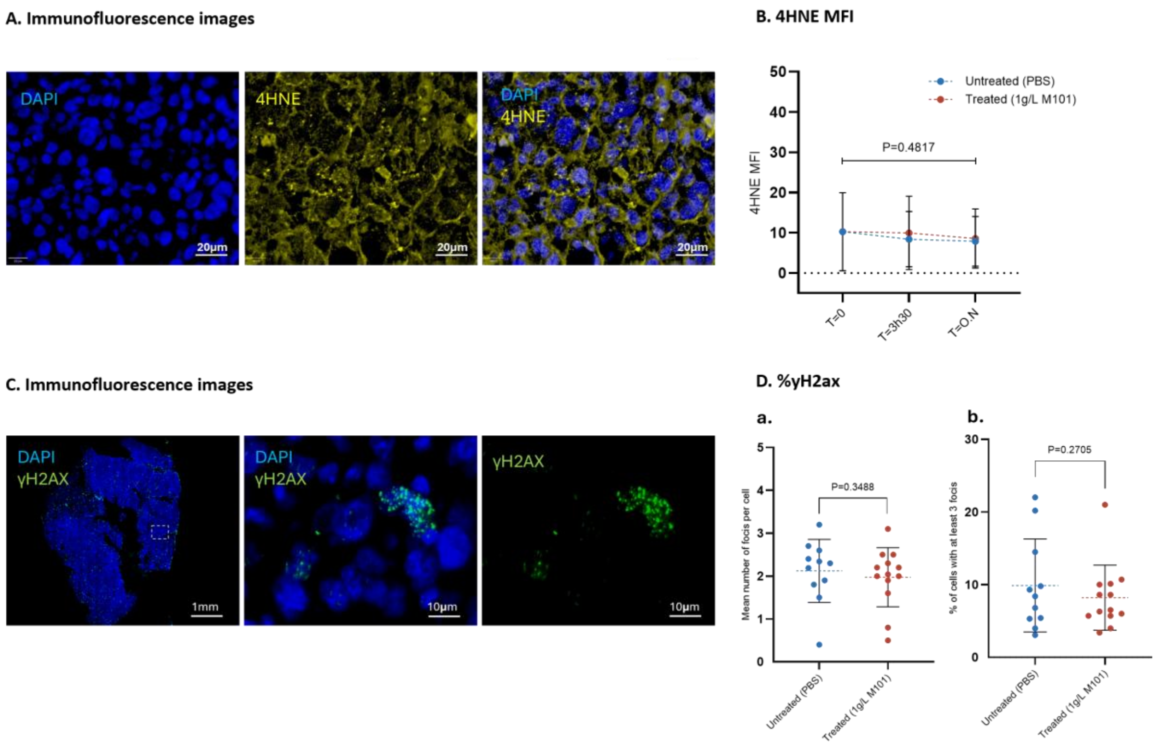


**Fig. S2 | Oxidative stress and DNA damage markers in preserved PC3 XG tissue.**

**A.** Representative immunofluorescence images of control (PBS) and M101-treated (1g/L) tissue stained for 4-HNE at T = 0, T = 3h30 and T = O.N. Scale bar, 20 µm. **B.** 4-HNE MFI over time. **C.** Representative immunofluorescence images of γ-H2AX staining in control and M101-treated tissue. Scale bar, 1 mm and 10 µm. **D.** Quantification of γ-H2AX as **(a)** mean number of foci per cell and **(b)** percentage of cells with at least three foci. Data are shown as mean ± SD, with each point representing one tumor, or the mean value for combined tumors. Statistical tests: Two-way ANOVA, exact P values are shown in the panels.
